# Touchscreen response precision is sensitive to the explore/exploit tradeoff

**DOI:** 10.1101/2024.10.23.619903

**Authors:** Dana Mueller, Erin Giglio, Cathy S. Chen, Aspen Holm, R. Becket Ebitz, Nicola M. Grissom

## Abstract

The explore/exploit tradeoff is a fundamental property of choice selection during reward-guided decision making. In perceptual decision making, higher certainty decisions are more motorically precise, even when the decision does not require motor accuracy. However, while we can parametrically control uncertainty in perceptual tasks, we do not know what variables - if any - shape motor precision and reflect subjective certainty during reward-guided decision making. Touchscreens are increasingly used across species to measure choice, but provide no tactile feedback on whether an action is precise or not, and therefore provide a valuable opportunity to determine whether actions differ in precision due to explore/exploit state, reward, or individual variables. We find all three of these factors exert independent drives towards increased precision. During exploit states, successive touches to the same choice are closer together than those made in an explore state, consistent with exploit states reflecting higher certainty and/or motor stereotypy in responding. However, exploit decisions might be expected to be rewarded more frequently than explore decisions. We find that exploit choice precision is increased independently of a separate increase in precision due to immediate past reward, suggesting multiple mechanisms regulating choice precision. Finally, we see evidence that male mice in general are less precise in their interactions with the touchscreen than females, even when exploiting a choice. These results suggest that as exploit behavior emerges in reward-guided decision making, individuals become more motorically precise reflecting increased certainty, even when decision choice does not require additional motor accuracy, but this is influenced by individual differences and prior reward. These data uncover the hidden potential for touchscreen tasks in any species to uncover the latent neural states that unite cognition and movement.

## Introduction

Sequential reward-guided decision making tasks, such as multi-armed bandit tasks, are well known to engage explore/exploit tradeoffs (Addicott et al., 2017; Chen et al., 2023a, 2021b; Ebitz et al., 2019, 2018; Stephens, 2008; Wyatt et al., 2023). Across species, exploration represents periods of variable choice selection and heightened learning about the environment, relative to exploit behaviors, which show consistent choice selection that is less sensitive to trial-to-trial feedback (Badre et al., 2012; Cavanagh et al., 2012; Daw et al., 2005; Frank and Fossella, 2011; Ting et al., 2023; Trudel et al., 2021). Explore/exploit tradeoffs therefore reveal that superficially similar choice behaviors (for example left vs right choice) can be driven by highly distinct neural states (Ebitz et al., 2020, 2019, 2018; Wang et al., 2023; Wyatt et al., 2023), likely reflecting differences in the certainty of choices. However, certainty in reward-guided tasks is individual and subjective, and we do not have good ways of measuring it without self report.

Perceptual decision making tasks reveal that higher certainty decisions are more motorically precise, even when the decision does not require motor accuracy (Follman et al., 2023; Palser et al., 2018; Sanchez et al., 2024; Wolpert and Landy, 2012). However, while we can parametrically control uncertainty in perceptual tasks, we do not know what variables--if any--shape motor precision during other forms of decision-making. This is an especially significant omission in the case of reward-guided decision-making because precision could be influenced either by prior rewards (which increase certainty about the correct action) or by decision-making states (which may or may not increase certainty). We do have reason to believe that explore/exploit states reflect differences in certainty - for example, exploitative choices are faster than exploratory ones (Addicott et al., 2017; Chen et al., 2023a, 2021a, 2021b; Ebitz et al., 2018; Hassall et al., 2013; Laureiro-Martínez et al., 2010; Walker et al., 2022; Wershbale and Pleskac, 2010). Although explore and exploit strategies are defined at the broadest level by the options chosen in a decision making task, these findings strongly imply that explore/exploit balance is also reflected in the fine-grained execution of the task.

Touchscreen operant chambers in animal models offer a powerful and novel approach in exploring the kinetics of a choice response by logging the precise coordinates and timing of each choice that is made on the screen, across thousands of choices. Screens by default offer no immediate, tactile feedback about choice accuracy, requiring that longer trial-and-error processes influence touch similarity. Pigeons and other birds have shown an awareness of spatial location of touches on touchscreens and make minute adjustments of touches as the task evolves, suggesting that the same might be evident for rodents (Capshew, 1993; Goodale, 1983; Jager and Zeigler, 1991; Peterson, 2004; Skinner, 1960; Spetch et al., 1992). We took advantage of this rich but underutilized data to analyze the location of decision touches across sexes from explore/exploit data in mice we have previously published (Chen et al., 2021b), asking if explore/exploit balance governed how similar choice touches were from one trial to the next. We found that actions become more precise in exploit state behavior compared to explore state. This effect was independent of a similar effect of reward on touch location, suggesting parallel mechanisms by which explore/exploit state and prior outcomes influence the precision of the next action execution. Because male and female mice employ different strategies in the two-arm restless bandit task, we tested whether the precision of choices to the screen was modulated by sex, and found that actions were more precise in females compared to males, also independent of the impact of explore/exploit state and reward experience, suggesting individual differences regulating action precision over and above other cognitive features of the task. Overall, this novel analysis capitalizes on the hidden potential for touchscreens to measure not only choice behaviors but the motor actions that generate them, informing the neural states that unite movement and cognition.

## Results

To understand how actions in the chamber are influenced by internal states in the animals and external events, we took advantage of a previously collected dataset examining sex differences in explore/exploit balance in mice in a touchscreen bandit task. Decision making data from the experiments analyzed here were originally shared in Chen et al. 2021. These data were collected from age-matched male and female wild-type mice (n = 32, 16 per sex, strain B6129SF1/J). Mice were trained in a two-arm spatial restless bandit task (**Figure 1a, 1c**) in a trapezoidal shaped touchscreen operant chamber. In this bandit task the probability of reward of each left and right choice changes independently and randomly of the other, with a 10% chance of probability change on each trial (**Figure 1c:** example probability walk). The unpredictability of this task encourages mice to continually learn and survey their choices, exploring to find the best option and exploiting a good rewarding option across a 300 trial session. Explore and exploit trials were labeled using a Hidden Markov model (HMM) approach (Chen et al., 2021b; Ebitz et al., 2018) where a mouse could either explore, exploit left choice, or exploit right choice (**Figure 1c**). Each trial nosepoke response on the touchscreen can therefore be identified as an explore or exploit choice (**Figure 1b**).

**Figure 1:**
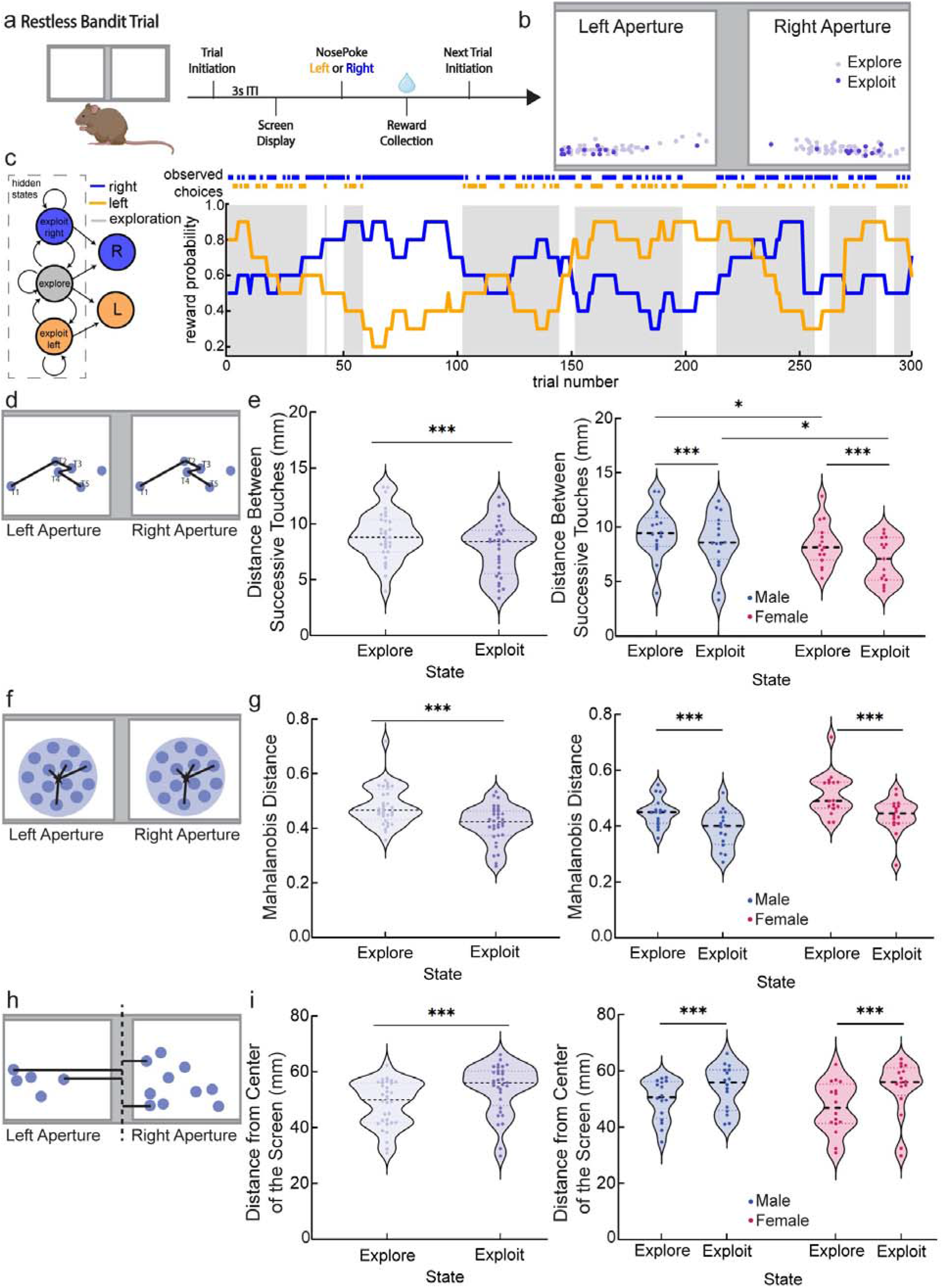
Exploit states and female sex reduce action variability during decision making. a) schematic depicting the timeline of a single trial. White squares indicate left/right spatial choice. b) An example of touch screen responses from one animal and one session, where light purple indicates explore touches and dark purple indicates exploit touches. c) Schematic depicting the hidden Markov model (HMM) and labeling explore trials along an example two-arm restless bandit probability walk. Orange traces indicate the probability and choices of left side touches. Blue traces indicate the probability and choices of right side touches. Gray shaded regions indicate HMM labeled explore trials. d) Schematic of Euclidean distance where the distance is calculated between touch 1 and touch 2, touch 2 and touch 3, touch 3 and touch 4, and so on. Shown here are possible left/right touches in blue and the distance relationship from one to another represented by black lines. e) Average Euclidean distance split by state (left) and sex (right). Exploit touches and females had significantly reduced Euclidean distance. Light purple indicates distance between explore touches and dark purple indicates distance between exploit touches. Red indicates female and blue indicates male mice. In violin graphs, individual data points are data from one mouse averaged across all sessions. f) Schematic of Mahalanobis distance where the individual data points are measured from the overall centroid of the dataset. Shown here are possible left/right Mahalanobis clusters (light blue circles) and centroids (stars) and the Mahalanobis distance relationship from each touch (darker blue circles) in a cluster to the centroid represented by black lines. g) Average Mahalanobis distance split by state (left) and sex (right). Exploit touches had significantly reduced Mahalanobis distance. Light purple indicates Mahalanobis distance between explore touches and dark purple indicates Mahalanobis distance between exploit touches. Red indicates female and blue indicates male mice. h) Schematic of distance from the center of the screen where touch distance from both left and right choice apertures is measured from the midpoint of the operant screen. Shown here are possible left/right touches in blue and the distance of each from the center of the touchscreen represented by black lines. i) Average distance from the center of the screen split by state (left) and sex (right). Explore touches were significantly closer to the center of the screen. Light purple indicates distance from the center of the screen for explore touches and dark purple indicates distance from the center of the screen for exploit touches. Red indicates female and blue indicates male mice. For simplicity of visualization, all plots are averages across trials and sessions, so that each individual data point plotted represents the overall average for a mouse. Significant throughout this paper is represented in the following way: * p value less than 0.05 and greater than 0.01; ** p value less than 0.01 and greater than or equal to 0.001; *** p value less than 0.001. Violin graphs depict median and quartiles of the dataset.

### Exploit states and female sex are associated with reduced action variability

Using previously assigned explore/exploit states for each trial, we examined the action associated with each choice, taking advantage of logging the coordinate locations of nosepokes in our touchscreen operant chambers. This allowed us to have a two dimensional location for each decision a mouse made across the entire touchscreen space. We started with an Euclidean analysis to quantify the distance between successive touch responses where T1 was compared to T2, T2 was compared to T3, T3 was compared to T4, so long as all touches were from the same choice aperture and state (**Figure 1d;** (Ebitz and Hayden, 2021; Walther et al., 2016)). One mouse was excluded from Euclidean analyses as they never had a sequence of choices on the same side in the same state consecutively. Distance between successive exploit touches was smaller and therefore less variable than successive explore touches (**Figure 1e**, GLM, main effect of state, p < 0.001). However, sex also played a role - female mice had shorter distances between successive touches than male mice (**Figure 1e**, GLM, main effect of sex, p = 0.01). These results suggest that exploit touches are more stereotyped and perhaps represent a more automated behavioral response than the same choice made during exploration, and suggests that these behaviors are more stereotyped overall in females than in males.

Although these data suggest that exploit choices are more stereotyped than exploration, Euclidean analysis can only compare distances between touches that are consecutively occurring on the same side, and in the same explore/exploit state. An alternative approach for calculating distance that permits all touches to remain in analysis is the Mahalanobis distance, a method for finding the distance between a point and the center of a distribution (**Figure 1f**) (Ebitz and Hayden, 2021; Walther et al., 2016). With Mahalanobis distance the entire cluster of data points was analyzed for each choice aperture, including both explore and exploit touches. We separated the population of touch responses into those happening in explore states and those in exploit states, and calculated separate Mahalanobis distances for exploit and explore touches from centroids within each left/right choice aperture, combining the data from both apertures across all trials and sessions and getting an average distance for each animal. The Mahalanobis distance of an average exploit touch from the centroid of all exploit touches was smaller and less variable than the distance of an average explore touch from the explore centroid (**Figure 1g**, GLM, main effect of state, p < 0.001). Unlike Euclidean analysis, we do not find significant sex differences in Mahalanobis distances (sex was dropped in the GLM model with the lowest AIC value). The difference between sex influences on Euclidean and Mahalanobis distances may reflect the trial-to-trial variability that Euclidean analysis captures versus the overall distribution captured by Mahalanobis analysis. However, both analyses reveal a main effect of explore/exploit state on touch variability - that exploit touches occur closer together in space with less variability than explore touches.

In maze tasks, as animals approach a choice point, they exhibit a behavior called vicarious trial and error (VTE) in which they move their head while surveying options to guide flexible decision making, that is reduced as choices become repetitive (George et al., 2023; Johnson and Redish, 2007; Redish, 2016; Tolman, 1948, 1939). This raised the possibility that in a touchscreen environment, flexible decision making may be reflected in the approach to the screen, allowing them to survey choices from a central location while exploring versus approaching directly towards one option when exploiting. To determine whether our mice might be exhibiting physical signs of deliberation between the left and right choice apertures during the explore state, we calculated the distance from the midpoint of the entire touchscreen between the two response apertures (**Figure 1h**). Explore touches happen significantly closer to the center of the screen, and thus closer to the opposite response aperture, than exploit touches (**Figure 1i**, GLM, main effect of state, p < 0.001). This did not differ by sex (GLM, no main effect of sex, p = 0.767). These results suggest that in an explore state mice exhibit a VTE-like behavior as they approach an area equidistant from both response apertures and deliberate between left and right choice. Conversely, in an exploit state, mice make responses committed to one aperture at a farther distance from the center of the screen.

### Previous reward is associated with reduced action variability separate from the effect of explore/exploit state

One potentially significant difference between explore and exploit states that might influence animal actions is a differing rate of reward across states. Exploit behavior is likely to result from prior success in obtaining reward, and thus exploit states might be expected to be associated with higher reward. Alternatively, reward may have a separate impact on action precision that is unrelated to explore/exploit state influences (Abe et al., 2011; Cashaback et al., 2017; Galea et al., 2015; Hasson et al., 2015; Izawa and Shadmehr, 2011; Nikooyan and Ahmed, 2015; Ramkumar et al., 2016; Therrien et al., 2016; Trommershäuser et al., 2003). To examine the impact of reward on touch location we separated trials by outcome: rewarded/not rewarded. To determine the impact of being rewarded on a previous trial, distance measurements were taken between one trial back (T_-1_) - labeled as *rewarded* or *non-rewarded* - and the current trial (T_0_). Euclidean and Mahalanobis distances for touches on trials following rewarded choices was smaller and less variable than those following non-rewarded touches (**Figure 2a**, GLM, main effect of reward, p < 0.001; **Figure 2c**, GLM, main effect of reward, p < 0.001). However, the effect of reward on action precision was independent of an effect of explore/exploit state on action precision, with both previous trial reward and explore/exploit state contributing main effects on the precision of choice responses (**Figure 2b**, GLM, main effect of reward, p < 0.001; **Figure 2d**, GLM, main effect of state, p < 0.001). Euclidean effects were stronger in females (**Figure 2b**, GLM, main effect of sex, p = 0.01 and a sex by state interaction **Figure 2b**, GLM, sex/state interaction, p = 0.039). As expected from prior Mahalanobis analysis, there was no influence of sex on Mahalanobis distances. These results suggest that while reward impacts touch location and minute adjustments in responding on the touchscreen, it does not overpower the state effects shown in **Figure 1**.

**Figure 2:**
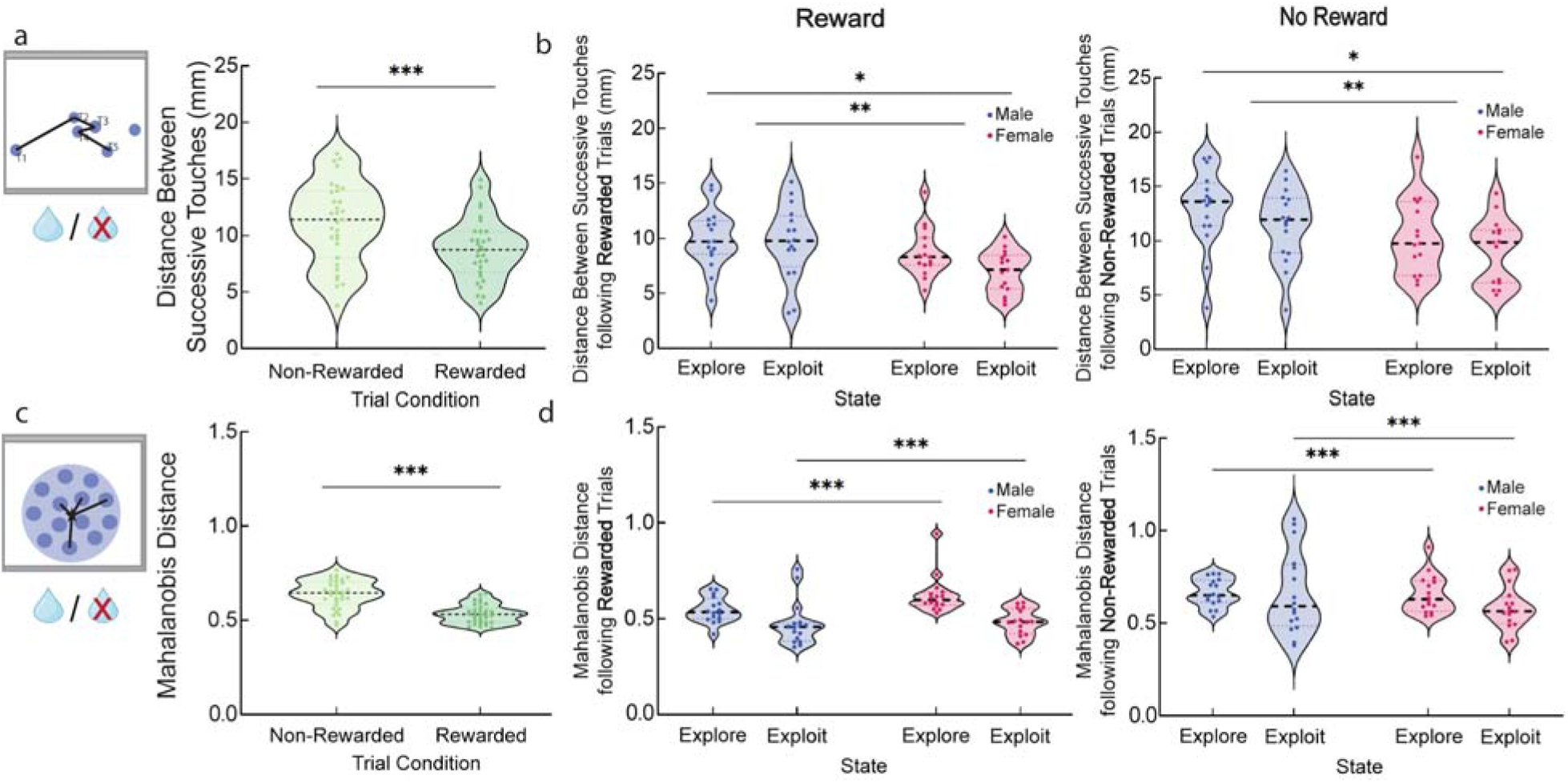
Previous reward reduces action variability independently from explore/exploit balance or female sex. a) Average Euclidean distance comparing rewarded vs. non-rewarded trials. Touches following rewarded trials had significantly reduced Euclidean distance. Light green indicates distance between non-rewarded touches and dark green indicates distance between rewarded touches. In violin graphs, individual data points are data from one mouse averaged across all sessions. b) Average Euclidean distance for rewarded (left) and non-rewarded (right) trials split by state and sex. Exploit touches and females had significantly reduced Euclidean distance. Red indicates female and blue indicates male mice. c) Average Mahalanobis distance comparing rewarded vs. non-rewarded trials. Touches following rewarded trials had significantly reduced Mahalanobis distance. Light green indicates Mahalanobis distance between non-rewarded touches and dark green indicates Mahalanobis distance between rewarded touches. d) Average Mahalanobis distance for rewarded (left) and non-rewarded (right) trials split by state and sex. Exploit touches had significantly reduced Mahalanobis distance. Red indicates female and blue indicates male mice. * p value less than 0.05 and greater than 0.01; ** p value less than 0.01 and greater than or equal to 0.001; *** p value less than 0.001. Violin graphs depict median and quartiles of the dataset.

In addition to modeling decision making behavior via Hidden Markov Models, we previously used reinforcement learning models to assess sex differences in latent parameters that could influence choice behavior, including learning rate parameter (alpha). We previously found in the animals in the current dataset that the alpha parameter was significantly higher in females, suggesting greater trial-to-trial influences of outcome on a female mouse’s next choice than on a male’s. Euclidean distance between touches is reduced by female sex, reward, and exploit behavior, and is a measure of trial-to-trial action variability. Therefore, we asked whether trial-to-trial action variability as measured by Euclidean distance between sequential touches on either aperture was correlated with trial-to-trial outcome sensitivity as measured by the alpha parameter for the best fit reinforcement learning model from (Chen et al., 2021b). With sex, distance, and alpha parameters as fixed effects, and individual mouse as a random effect, the GLM revealed a higher alpha parameter, indicating more rapid outcome sensitivity/value updating/learning rate, was associated with smaller distances between successive touches (GLM, main effect of alpha, p = 0.046), suggesting that animals that were more sensitive to outcomes in their choice behavior as measured by a reinforcement learning model also show greater precision of their actions. Additionally, we replicated the sex difference in touch precision with females having shorter distances (GLM, main effect of sex, p = 0.018).

### Exploit states and female sex reduce centroid shifting across session

Given the difference between sex influences and the consistency of state influences on Euclidean (**Figure 1e**) and Mahalanobis (**Figure 1g**) distances, we wanted to determine if the pattern of responding shifts differently across a session for male/female mice and explore/exploit state. Given that during exploration, animals are more likely to make choices closer to the midpoint of the screen (**Figure 1j**), it could be the case that exploration can be seen in terms of not only which aperture is chosen, but what section of the aperture responses in an explore state center on. Although we observe animals sampling between two response locations (right and left) in our decision task, it is not clear whether animals are behaving as though they are sampling two discrete options versus sampling touching an area in space. To measure this we separated each session into state *bouts*. A *bout* is defined as a period of touches within one state on a particular choice aperture. State transition trials from either explore to exploit or exploit to explore trigger a new *bout*. To determine how our mice use the available space within the choice aperture we calculated area and perimeter associated with each state bout. Bouts of touches were plotted and overlaid onto 2D contour plots from Plotly Graphing Libraries (**Figure 3a**). For each bout, Open Source Computer Vision (OpenCV) was used to capture the contours (bin traces) along continuous boundaries of the contour plots and calculate area and perimeter for the outermost bin - which is recognized as the outer range of nosepoke responses.

**Figure 3:**
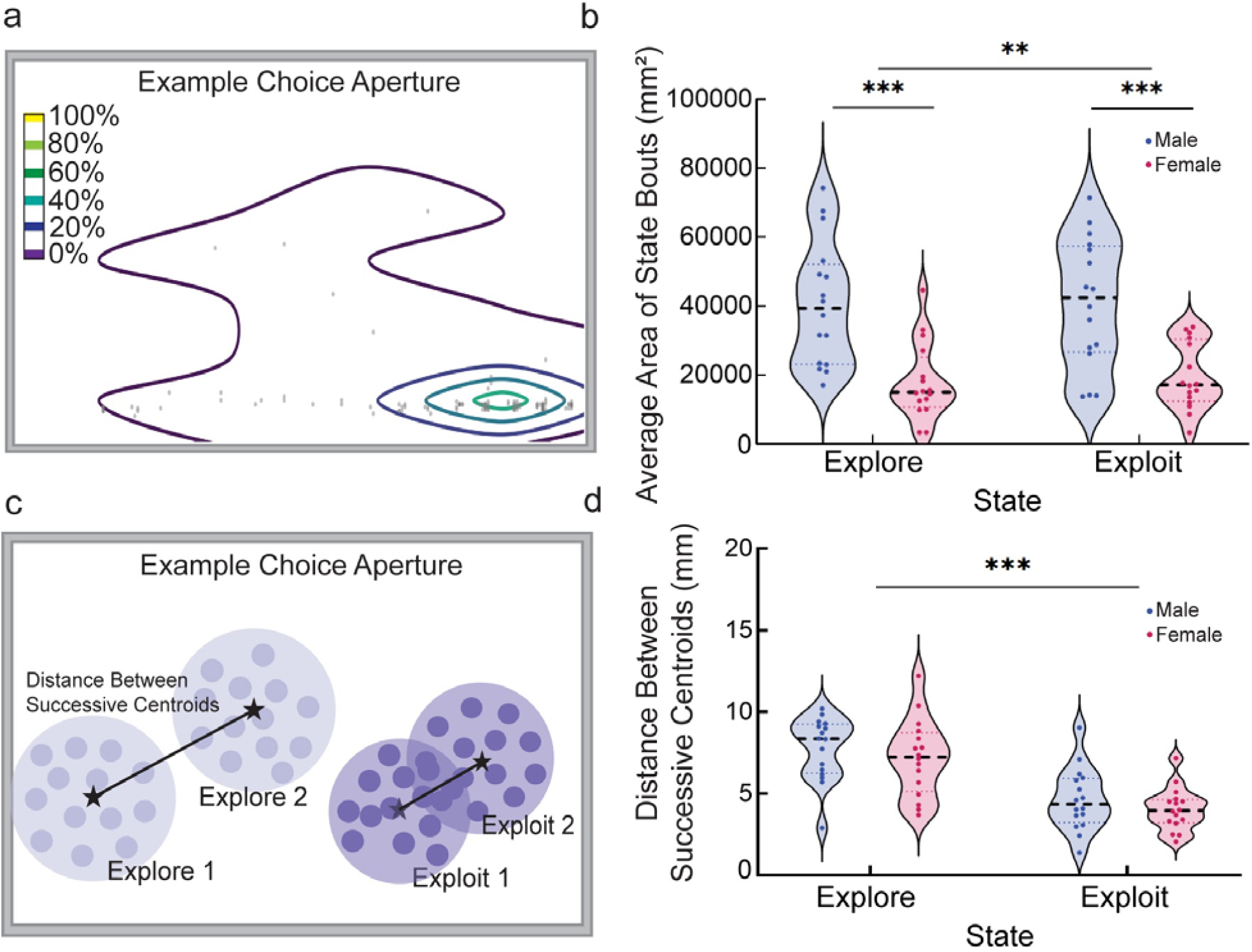
Exploit states and female sex reduce the total response space chosen across a decision making session. a) An example 2D contour plot from Plotly Graphing Libraries fit over our nosepoke touch locations to visualize the density and range of choice responding. Small gray circles are nosepoke touches within the bout of response data. Color map corresponds with density of data points within each bin, where the darkest purple (outer bin) is the least dense contour bin, which is used to calculate area and perimeter of the bout. b) Average area of bouts split by state and sex. Exploit touches and females had significantly reduced area. Red indicates female and blue indicates male mice. In violin graphs, individual data points are data from one mouse averaged across all sessions. c) Schematic depicting centroid shifts, where the Euclidean distance between two successive Mahalanobis centroids is calculated. Stars represent example centroids associated with bouts and black lines represent the distance calculations between those centroids. d) Centroid shifts split by state and sex. Centroid shifts were significantly smaller for exploit bouts. Red indicates female and blue indicates male mice. * p value less than 0.05 and greater than 0.01; ** p value less than 0.01 and greater than or equal to 0.001; *** p value less than 0.001. Violin graphs depict median and quartiles of the dataset.

Regarding the area of the touchscreen choice apertures used by the mice, exploit bouts occupied a smaller area (mm^2^) on the screen and were less variable than explore bouts (**Figure 3b**, GLM, main effect of state, p = 0.006). Female mice used significantly less area of the screen per bout than males (**Figure 3b**, GLM, main effect of sex, p < 0.001). The model used included an interaction term between state and sex, which was not significant (**Figure 3b**, GLM, interaction state/sex, p = 0.989). Perimeter of the touchscreen choice apertures used by the mice, exploit bouts occupied a smaller boundary (mm) on the screen and were less variable than explore bouts (GLM, main effect of state, p < 0.001). Female mice occupied a smaller boundary on the screen and were less variable than bouts by male mice (GLM, main effect of sex, p = 0.004). The model used included an interaction term between state and sex, which was not significant (GLM, interaction state/sex, p = 0.168). Further suggesting differences in touchscreen navigation across state and sex, where exploration and males interact with more overall area of the screen.

Each new bout of responding includes its own centroid, and these centroids may minutely move across the screen throughout a session, adjusting based on past experience. In a combination of analysis techniques, **Figure 3c** shows how the distance between successive centroids is calculated using the x,y centroid coordinates - as determined by the Mahalanobis analysis. Distances between centroids for successive exploit bouts were smaller and less variable than distances between centroids for successive explore bouts (**Figure 3d**, GLM, main effect of state, p < 0.001). We found that touches occurring during one bout of exploration were farther and more variable in distance from other bouts of exploration compared to more similar touch patterns across bouts of exploitation. Given that mice are using more overall screen space during explore than exploit trials, this further increases the likelihood that mice may be exploring individual touch locations over and above sampling just the left/right options we define.

## Discussion

The explore/exploit tradeoff is a fundamental property of choice selection during reward-guided decision making. Explore and exploit states are mediated by distinct neural circuit activity and reflect slower versus faster decision processes (Ebitz et al., 2020, 2019, 2018; Wang et al., 2023; Wyatt et al., 2023) and so likely reflect different levels of subjective certainty in a choice. Here, we take advantage of the observation that higher certainty actions in perceptual tasks are more precise to ask whether exploit states, reward feedback, or other factors lead to increased precision of choices. Using touchscreen operant chambers in mice we asked whether explore/exploit balance governed the precision of actions during decision making, finding independent effects of (1) explore/exploit state, (2) prior reward, and (3) sex on increasing similarity of touches. These data suggest multiple independent mechanisms regulate the precision of actions associated with choices and that the explore/exploit state is visible at the level of motor performance.

Perceptual decision making tasks reveal that higher certainty decisions are more motorically precise, even when the decision does not require motor accuracy (Follman et al., 2023; Palser et al., 2018; Sanchez et al., 2024; Wolpert and Landy, 2012). One striking result from our study is that exploit states reduce action variability in choice location. This suggests that exploit touches reflect higher certainty in the animal. The similarity of touch locations suggests that exploit decisions are more repetitive, stereotyped, or automated behavioral responses (Dezfouli and Balleine, 2012; Dolan and Dayan, 2013; Gillan et al., 2016; Yin et al., 2004). Exploit choices happen faster in comparison to explore choices (Chen et al., 2023b, 2021c; Ebitz et al., 2018), an expression of these cognitive strategies at the motor level (Carsten et al., 2023; Chen et al., 2017). Stereotyped performance of a behavior has previously been linked to a lack of deliberation (Foster, 1998; Graybiel, 2008; Mitchell and Etches, 1977; Smith and Graybiel, 2016). Our findings are broadly consistent with the idea that exploit choices reflect behavioral automation of a higher confidence response, while explore reflects deliberation.

Exploration and deliberation processes involve the subject surveying options (Gilbert and Wilson, 2007; Payne et al., 1993; Rangel et al., 2008). Deliberation is physically expressed through pausing, slower decision making, and “vicarious trial and error” behavior, reflecting forward thinking and prospective deliberation (Dolan and Dayan, 2013; George et al., 2023; Johnson and Redish, 2007; Redish, 2016; Tolman, 1948, 1939). We observed that explore touches happen significantly closer to the center of the screen than exploit touches, which implies animals are approaching exploratory choices between the two apertures, rather than from off to one side. In addition, we found that touches occurring during one “bout” of exploration were farther from other bouts of exploration compared to exploit. Given that mice are using more overall screen space during explore than exploit trials, this suggests mice may be exploring individual touch locations across the screen over and above sampling just the left/right options we define. Self-directed exploration may reflect an increasingly fine-grained goal-directed search for the most rewarding action, similar to autoshaping.

A potential confound between explore/exploit state and action precision is that exploit actions are more likely to be reinforced. However, exploit states and prior reward independently reduced action variability. This suggests that while reward may cause trial-to-trial adjustments in responding on the touchscreen, reward does not overpower the state effect. Reward-triggered changes in response precision may be a function of individual reward sensitivity. Animals with a higher learning rate derived from a reinforcement learning model showed smaller distances between successive touches, suggesting that reward sensitivity varying across individuals is associated with increased action precision. This effect was larger in females than in males, highlighting sex as a third independent factor governing choice precision.

The data in this manuscript were previously used to reveal a sex difference in the balance of explore/exploit strategies (Chen et al., 2021c). Because male and female mice employ different strategies in the two-arm restless bandit task, we sought to test whether motor responses associated with the different strategies were physically different in distribution and spatial location. We found that actions were more precise in females compared to males, independent of the impact of explore/exploit state and reward experience, suggesting individual differences regulating action precision over and above moment to moment features of the task. However, not all explore/exploit differences were sex different. In particular, there was no sex difference in how close animal responses were to the center of the screen during exploration. This suggests that the overall deliberative process of an exploratory decision is probably similar across sexes, but the sequential execution of these decisions are more similar in females than males. Overall these findings agree with a growing literature that finds male decision and/or motor behavior to be more variable than females in rodents (Chen et al., 2021a; Levy et al., 2023) and humans (Dosenbach et al., 2017).

Touchscreens are increasingly used not only by rodent researchers, but by people working with humans via smartphone-mediated ecological assessments. Our analysis reveals a powerful way to evaluate the distribution and consistency of motor behaviors in choice responding. Motor abnormalities are a common feature across patients with psychosis (Walther and Mittal, 2017), autism (Mody et al., 2017; Mosconi and Sweeney, 2015), and depression (Sobin and Sackeim, 1997), and explore/exploit tradeoffs reveal neuropsychiatric influences (Addicott et al., 2017; Wyatt et al., 2023). The increasing prevalence of touchscreen phone testing in human neuropsychiatric research raises the distinct possibility of analyses of touch responses (Azenkot and Zhai, 2012; Gosling and Mason, 2015; Harari et al., 2016; Intarasirisawat et al., 2019; Miller, 2012) as a novel cross-species translational measure of explore/exploit tradeoffs.

## Methods

### Subjects

Animals were thirty-two 129/B6J F1 mice (16 male and 16 female) from The Jackson Laboratory. Behavioral data from these mice running this task were previously published by the lab (Chen et al., 2021b). Colony rooms were temperature controlled (20.5°C; 69°F) and on a light-dark cycle of 12 hours with the lights off at 9am. Mice were housed in groups of four with water ad libitum. Mice were food restricted to no lower than 85% of their free-feeding body weight. All animals were cared for according to the guidelines of the National Institution of Health and the University of Minnesota (UMN) and UMN IACUC approval.

### Behavioral Data

Details on the methods for behavioral training and the restless bandit task are published in (Chen et al., 2021b). Behavioral testing was carried out in the same touchscreen chambers for all mice throughout the present study (Lafayette Instrument Company, Lafayette, IN). Computational models were fit to mouse data in this paper, including a hidden Markov model (HMM) and an RLCK reinforcement learning model (Chen et al., 2021b). The HMM was used to determine when animals were exploring or exploiting their options in the restless bandit task, where P(exploration) is the probability of mouse exploration between choices. The previous manuscript compared several different RL models and identified the strongest fit to animal behavior from an RLCK model, which captures both value-based and value-independent decisions using the following four parameters: learning rate, decision noise, choice bias, and choice stickiness. Here we use this RLCK model’s alpha parameter compared to distance between successive touches to assess how learning rate impacts micro adjustments to spatial touch locations across sex. For validation of both models please see (Chen et al., 2021b) eLife publication.

### Coordinate Analysis

The Bussey-Saksida touchscreen apparatus (Lafayette Instrument Company) is sensitive to continuous and rapidly repeated touches in the same location and across the entirety of the screen (Heath et al., 2015). Each touchscreen represents the x,y coordinates of each response an animal makes on the screen from IR beam technology where IR emitters are positioned along two sides of the screen (i.e. top and right sides) and IR receivers are positioned along the other two sides of the screen (i.e. bottom and left sides). In this configuration, IR beams are ideally suited to determine the shadow of the touch to triangulate the location of choice response. IR beam configuration results in a touch resolution that matches the monitor resolution of 800×600 pixels. **Figure 1b** visualizes this data, representing the choices of four different mice selecting between two options on the touchscreen over 300 trials, with explore responses in the lighter purple and exploit responses in the darker purple. **Figure 1b** provides an example of nosepoke responses for one mouse across a session and the change in touch pattern between explore/exploit touches as identified by our HMM. Left and right touchscreen choice apertures are 240×240 pixels each, never change position or size, and x,y coordinates are separately generated for each touch aperture. Throughout all analyses we have transformed pixels into millimeters. 1 pixel is 0.29 millimeters. Unless mentioned otherwise, for all data, a GLM stepwise model selection analysis was used to determine the optimal model with the lowest AIC value and p values are shared from those most optimal models.

### Distance from the Center of the Screen

The spatial split in exploration and exploitation visualized by these plots suggested that explore trials were closer to the center of the touchscreen than exploit trials were, prompting us to quantify the distances (**Figure 1i**). With the center of the screen being 400 out of 800 total pixels (width of the screen), the difference between the x pixel coordinate of the x,y location of each touch response and 400 pixels was calculated and converted into millimeters. An absolute value is applied so that the distance away from the center of the screen is always a positive value to reflect distance. This calculation was done across all touches in every session. Trials were split by explore and exploit and all data was averaged across all eight restless bandit sessions for graphing purposes.

> Example (x,y) is (34,208).
>
> Distance from the center of the screen = |400 - x|
>
> Distance from the center of the screen = |400 - 34| = 366 pixels.

### Euclidean Analysis

The first method we used to quantify the distance between nosepoke touches was a Euclidean Analysis (Ebitz and Hayden, 2021; Walther et al., 2016) in which we used the pythagorean theorem to calculate the hypotenuse between two points with (x,y) coordinates that were successive, from the same choice aperture (left/right), and within the same HMM decision state (explore/exploit) (**Figure 1d**). In python this calculation was done using numpy.hypot(). A drawback of this analysis is the amount of data points that get excluded given that the included data points must be consecutively from the same choice aperture side and within the same state. Distances were split by explore and exploit and all data was averaged across all eight restless bandit sessions for graphing purposes. In the example below “T” represents touch (nosepoke).

> Example T_1_ is (x_1_,y_1_) and T_2_ is (x_2_,y_2_).
>
> Distance Between Successive Touches (hypotenuse) = 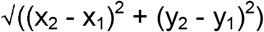

### Mahalanobis Analysis

The second method we used to quantify touch patterns was a Mahalanobis analysis (Ebitz and Hayden, 2021; Walther et al., 2016) where, unlike the Euclidean analysis, we didn’t have to exclude any touch data points. With this analysis we were able to calculate separate centroids based on the data clusters for both the left side touches and right side touches and calculate the distance of each touch coordinate from each overall centroid (**Figure 1f**). The centroid is the central point in the data field that can be considered the overall mean for multivariate data given that this is the point where all means from all variables intersect. The further away a data point (touch) is from the centroid, the larger the Mahalanobis distance value. Distances were split by explore and exploit and all data was averaged across all eight restless bandit sessions for graphing purposes. In the formula below X_A_ and X_B_ represent a pair of objects, which are the x and y coordinates; C is the sample covariance matrix, calculated using numpy.cov() in python; and T is the transposition of the matrix over its diagonal, calculated using numpy.linalg.inv() in python.

> Mahalanobis Distance = [(X_B_ - X_A_)^T^* C^-1^* (X_B_-X_A_)]^0.5^

### Reward

To determine whether being rewarded in the restless bandit task impacts touch location, we compared trial outcome (rewarded or non-rewarded) from the previous trial (T_-1_) to the change in touch location on the current trial (T_0_). This was done using both Euclidean and Mahalanobis analyses.

### Distance Between Successive Bouts

To understand how touches were organized within and across periods of exploration or exploit as defined by HMM, we divided the data into “bouts”. Rather than looking at our nosepoke data clusters throughout an entire session, a “bout” is described as a period of touches within one HMM defined behavioral state on one particular choice aperture. Thus, explore states may contain separate bouts on the left or right side, but these are analyzed separately. State transition trials from either explore to exploit or exploit to explore trigger a new “bout.” By looking at individual state bouts of choice responding, we can investigate whether explore or exploit centroids on a given response area are shifting more throughout a session. This analysis combines both Euclidean and Mahalanobis methods previously described. Mahalanobis analysis is used to determine the centroid of each individual “bout.” From here, the distance between successive centroids is calculated using the Euclidean analysis, which employs the pythagorean theorem (**Figure 3c**). Distances were split by explore and exploit and all data was averaged across all eight restless bandit sessions for graphing purposes. In the example below “C” represents centroid.

> Example C_1_ is (x_1_,y_1_) and C_2_ is (x_2_,y_2_).
>
> Distance Between Successive Touches (hypotenuse) = 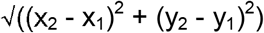

### Contour Plots and Area Calculations

In order to calculate the amount of space occupied by each bout we calculated the area and perimeter of the bouts. In Python, 2D contour plots from Plotly Graphing Libraries were fit over our nosepoke touch locations to visualize the density and range of choice responding. Bins edges were designated by numpy.histogram and filtered at every-other bin so they were twice as big as the standard output. The color bar was fixed from 0 to 1 across all generated plots to ensure consistency of calculations **(Figure 3a)**. Contour fill was removed, leaving just the outlines at a thickness of “3” so the trace would be better recognized by OpenCV.

Once a contour plot was generated for each bout, Open Source Computer Vision (OpenCV) was used to capture the contours along continuous boundaries and calculate area (cv.contourArea) and perimeter (cv.arcLength) for each bin. While tracing the contours, cv.threshold was set to cv.THRESH_BINARY and cv.findContours was set to cv.CHAIN_APPROX_SIMPLE. Contour Approximation was used when it was necessary to approximate the area between two separate contour groups. We focused on the dimensions of the outermost bin as the best representation for the spread of data throughout a bout **(Figure 3a)**. The outermost bin was filtered using the structure hierarchy, or rather the nested orientation of the contours labeled numerically with “parent” and “child” identifications. Areas and perimeters of bouts were split by explore and exploit and all data was averaged across all eight restless bandit sessions for graphing purposes.Finally, area and perimeter were calculated for the correctly identified contour bin. OpenCV was run through Minnesota Supercomputing Institute (MSI).

## Acknowledgments

This work was supported by NIMH R01 MH123661, NIMH P50 MH119569, Canada Research Chair CRC-2022-00192 (RBE), NSERC RGPIN-2020-05577 (RBE), NIMH T32 training grant MH115886, NIDA T32 training grant DA050560, and NIDA T32 training grant DA007234. We thank Nic Glewwe for comments on an earlier draft of the manuscript and Matt Croxall from Lafayette Instrument Company for valuable feedback and technical support.

## Notes

### Competing Interest Statement

The authors have declared no competing interest.

